# Sex-Dimorphic Aging of Cardiovascular Disease Genes: A Network-Based Multi-Omics Analysis

**DOI:** 10.64898/2026.07.08.737220

**Authors:** Annamaria Defilippo, Fabiola Boccuto, Pietro Hiram Guzzi, Pierangelo Veltri

## Abstract

Sex differences influence the incidence, timing, clinical presentation, and outcomes of cardiovascular disease (CVD), yet the molecular programs through which aging interacts with biological sex remain insufficiently understood. To address this gap, we integrated basal gene expression profiles from multiomics data across 981 donors and 17 CVD-relevant tissues with regulatory, genetic, network, disease-expression, and druggability information to characterize sex-dimorphic aging patterns in 1,176 candidate CVD genes.

Using a two-step expression analysis, we identified 4,404 genes with significant age-associated expression trends (BH-FDR *<* 0.05), including 2,718 male-specific, 202 female-specific, and 742 shared trends. Concordant evidence across complementary statistical approaches highlighted 35 high-confidence sex-dimorphic genes, including *REN*, *APOE*, *GUCY1A2*, and *SRD5A2*. Regulatory analysis showed that most CVD genes were influenced by nearby genetic variants, with 96.2

Network-based analyses further suggested that CVD genes are organized within hierarchical biological structures, with curated protein-interaction data showing stronger geometric organization than broader interaction resources. Integration with Open Targets identified 289 genes already linked to approved drugs and 48 of the top 50 biomarker candidates supported by GWAS–eQTL colocalisation evidence. A final composite ranking prioritized *NTRK1*, *TUBB4A*, *PTGS2*, *IL6*, and *PDE5A*, and identified 19 actionable biomarkers supported by convergent expression, regulatory, genetic, and therapeutic evidence. Among these, a dedicated sex-specific evidence score nominated *GUCY1A2*, *CACNA1D*, *PGR*, *PDE5A*, and *LEPR* as the strongest candidates for sex-stratified validation, with *GUCY1A2* and *PDE5A* converging on a nitric oxide–cGMP signaling axis.

This study provides an integrative framework for discovering sex-dependent molecular signatures of cardiovascular aging and for prioritizing biologically supported, potentially actionable targets for precision cardiovascular medicine.

## 1 Introduction

Cardiovascular disease (CVD) remains the leading cause of death worldwide and shows marked differences between men and women in disease onset, clinical phenotype, progression and outcome. Men generally develop CVD earlier in life, whereas women more frequently present with conditions such as heart failure with preserved ejection fraction, coronary microvascular dysfunction and spontaneous coronary artery dissection. These clinical differences have often been interpreted through the lens of sex hormones, but increasing evidence indicates that biological sex also acts through broader molecular mechanisms, including sex-biased gene expression, X-chromosome escape, tissue-specific regulation and sex-dependent transcriptional programs [1, 2, 3].

Despite this progress, the interaction between sex and aging remains incompletely understood at the molecular level [4, 5]. Aging is the dominant risk factor for most cardiovascular conditions, but age-related molecular trajectories may not be identical in men and women. Identifying genes whose expression changes with age in a sex-specific manner is therefore essential to understand why cardiovascular risk, disease phenotype, and treatment response diverge across the life course. This information may also help move the field beyond demographic sex as a simple covariate toward a more mechanistic view of sex-aware cardiovascular biology.

Public existing databases offer a valuable data source to investigate this question across human tissues [6]. For instance, the Genotype-Tissue Expression (GTEx) [6] includes gene expression data from 981 donors across 49 tissues, together with information on sex, age and tissue source. Previous large-scale GTEx analyses have shown that more than one third of genes display sex-biased expression in at least one tissue, and that sex differences also extend to regulatory and transcription factor networks [1, 2]. However, to the best of our knowledge there is still room for other studies focusing on CVD genes or examined aging as a sex-interacting variable across cardiovascular and metabolically relevant tissues.

From a translational perspective, sex-dependent molecular signatures may complement conventional cardiovascular risk factors and circulating biomarkers [7]. They may support more precise patient stratification, identify sex-specific disease mechanisms and reveal candidate targets whose biological relevance depends on tissue context and age. These goals require an integrative approach, because expression changes alone cannot establish whether a candidate gene is genetically regulated, embedded in relevant interaction networks or potentially druggable [8].

We previously showed that age-related expression trends in type 2 diabetes comorbidity genes can be systematically characterized using GTEx data and non-parametric age-trend testing [9]. Here, we extend that framework to cardiovascular disease by explicitly incorporating biological sex. We combine sex-stratified age-trend analysis with interaction modelling to distinguish genes that show age-associated changes in one sex from those with formal sex-by-age effects.

To connect these expression signatures to regulatory and functional context, we further integrate cis-eQTL evidence, SuSiE fine-mapping, protein interaction networks and network geometry. Protein interaction resources such as BioGRID [10] and STRING [11] provide complementary views of the CVD interactome: one emphasizing curated physical interactions, the other integrating multiple evidence channels. Network geometry, including Ricci curvature and hyperbolic embedding, adds a structural perspective by identifying hub regions, bridging edges and hierarchical organization that are not fully captured by standard degree-based measures [12, 13, 14, 15].

Finally, we connect molecular discovery to putative therapeutic interpretation by incorporating Open Targets evidence, druggability and GWAS–eQTL colocalisation [16, 17, 18]. This allows candidate genes to be prioritized not only by expression differences, but also by genetic support, network position and pharmacological tractability. The resulting framework is designed to identify robust, interpretable and clinically relevant molecular signatures of sex-dimorphic cardiovascular aging.

## 2 Methods

### 2.1 Gene Set Curation and GTEx v11 Data Acquisition

We assembled two complementary CVD gene panels. Panel A was derived from the T2DiACoD framework selecting 159 genes with relevance to cardiovascular comorbidity. Panel B was curated from the Open Targets Platform [16] by querying six major CVD entities: coronary artery disease, heart failure, atrial fibrillation, hypertension, myocardial infarction and atherosclerosis. Targets with an overall association score of at least 0.4 were retained. The union of the two panels yielded 1,176 unique candidate CVD genes, with 48 genes shared by both panels (Jaccard index = 0.041).

Raw gene-level read counts were obtained from GTEx v11 for 17 tissues selected for cardiovascular, metabolic or vascular relevance: heart atrial appendage, heart left ventricle, aorta, coronary artery, tibial artery, liver, subcutaneous adipose tissue, visceral adipose tissue, pancreas, kidney cortex, whole blood, lung, skeletal muscle, tibial nerve, cortex, hippocampus and hypothalamus. Subject-level sex and age-bin information, together with sample annotations, were retrieved from the corresponding GTEx metadata. The final dataset included 981 subjects (654 male and 327 female) and 8,050 CVD-filtered samples across the selected tissues.

### 2.2 Expression Analysis: Kruskal–Wallis, DESeq2, and Disease-State Validation

For each gene–tissue pair, age-associated expression trends were tested separately in male and female samples using Kruskal–Wallis tests across six age bins. Age–sex strata with fewer than five samples were excluded, and only tests with at least three valid age groups were retained. Multiple testing corrections were performed using both Bonferroni and Benjamini–Hochberg false-discovery rate (BH-FDR) procedures in all 38,694 tests. The trend direction was summarized using the Spearman correlation between the median expression and the age-bin midpoint, with positive, negative, or non-monotonic trends assigned according to the correlation value. Genes were then classified as male-specific, female-specific or shared according to the pattern of significant sex-stratified trends.

To formally test whether age effects differed by sex, DESeq2 [19] was applied independently to each tissue using the design formula ∼ SEX+AGE_centered_ +SEX : AGE_centered_. The centered age term was computed from the age-bin midpoint and scaled to zero mean. This model produced three coefficients per gene: age main effect, sex main effect and sex-by-age interaction. Log fold-change shrinkage was performed using apeglm [20], and significance was defined as padj *<* 0.05. In total, 59,808 coefficient-level tests were performed across 17 tissues.

To assess whether basal age- and sex-related signatures were also altered in disease, two independent GEO datasets were analysed: GSE116250, comparing failing and non-failing left ventricular heart samples, and GSE55296, comparing ischemic or dilated cardiomyopathy with control heart samples. Because both datasets provided normalized expression values, differential expression was assessed using limma-voom [21]. Genes were considered differentially expressed when adj.P.Val *<* 0.05 and | log_2_ FC| ≥ 1.

### 2.3 Network Construction and Ricci Curvature

Protein interaction networks were built by restricting BioGRID [10] and STRING v12.0 [11] to the 1,176 candidate CVD genes. BioGRID was limited to curated human physical interactions, resulting in a network of 1,023 nodes and 9,473 edges. STRING was queried for human interactions at a medium confidence threshold and yielded 1,093 nodes and 9,311 edges. STRING enrichment analysis was used to summarize pathway and biological process content within the network.

Ricci curvature was computed on the largest connected components of both networks using the GraphRicciCurvature package. Ollivier–Ricci curvature [12] quantifies the transport distance between the neighborhoods of adjacent nodes; negative values typically identify bridge-like edges connecting distinct network regions, whereas positive values identify locally clustered edges. Forman–Ricci curvature [13] provides a complementary discrete approximation based on local edge and node structure. Sex-stratified subnetworks were generated from Kruskal–Wallis classifications, and curvature metrics were recomputed on the induced male-specific, female-specific and shared subnetworks. Differences between curvature distributions were tested using Mann–Whitney *U* tests.

### 2.4 Network Geometry: Topology, Hyperbolicity, and Embedding

To characterize network organization beyond local curvature, we computed standard topological measures on the largest connected components, including degree assortativity [22], Louvain modularity [23], small-world index [24], rich-club coefficient [25], diameter, eccentricity and *k*-core structure. We also computed a node-level combinatorial Gaussian curvature, *κ*(*v*) = 1 − deg(*v*)*/*2 + tri(*v*)*/*3, where tri(*v*) is the number of triangles involving node *v*. Associations between curvature and topology were assessed using Spearman correlations between edge curvature and edge betweenness, and between node curvature, degree and betweenness.

Global hyperbolicity was estimated using the Gromov four-point condition [26]. For each random quadruple of nodes (*a, b, c, d*), we computed *δ* = (max(*s*_1_*, s*_2_*, s*_3_) − median(*s*_1_*, s*_2_*, s*_3_))*/*2, where *s*_1_ = *d*(*a, b*) + *d*(*c, d*), *s*_2_ = *d*(*a, c*) + *d*(*b, d*) and *s*_3_ = *d*(*a, d*) + *d*(*b, c*). We sampled 50,000 quadruples for each network and compared the resulting values with configuration-model null networks that preserved the degree sequence.

Poincaré disk embedding was used to evaluate whether graph distances were better represented in hyperbolic than Euclidean space [14, 27]. Radial coordinates were assigned from degree or eccentricity, placing highly connected nodes closer to the disk center. Angular coordinates were derived from graph Laplacian eigenvectors. Embedding quality was measured as the Spearman correlation between embedded distances and shortest-path distances, and compared with a two-dimensional Euclidean spectral embedding. The same procedure was applied to sex-stratified subnetworks.

### 2.5 eQTL Analysis: Tissue Specificity and Fine-Mapping

GTEx v11 single-tissue cis-eQTL summary statistics and SuSiE fine-mapped credible sets were used to characterize regulatory support for the CVD gene set. Per-tissue eGenes were filtered at qval ≤ 0.05 and matched to candidate genes using gene symbols. Fine-mapped variants were retained when their posterior inclusion probability (PIP) exceeded 0.5.

For each gene, we summarized eQTL evidence across tissues by extracting the strongest eQTL per tissue, its effect size and its direction. Tissue specificity was quantified using *τ* = 1 − (mean|slope|*/*max|slope|), where higher values indicate more tissue-restricted regulation. We also assessed whether eQTL effects had concordant direction across tissues, clustered tissues by shared eGenes and summarized SuSiE credible set size, maximum PIP and cross-tissue recurrence of high-PIP variants. Finally, expression and regulatory layers were integrated by testing whether eQTL presence, eQTL tissue breadth and high-PIP fine-mapping evidence were associated with Kruskal–Wallis, DESeq2 and overall evidence counts.

### 2.6 Open Targets Integration and Biomarker Ranking

Open Targets evidence was used to place candidate genes in a disease and therapeutic context [16]. Disease association scores and datatype-specific evidence were retrieved for coronary artery disease and heart failure, and filtered to the CVD gene set. For each gene, we also collected information on drug and clinical candidates, tractability, Reactome pathways [28] and safety liabilities. Tractability was summarized into four tiers: approved drug, clinical-stage candidate, preclinical or structural support, and no available tractability evidence. For the top biomarker candidates, GWAS credible sets, L2G predictions [29] and colocalisation evidence [17] were used to evaluate whether genetic association and molecular QTL signals likely shared the same causal variant.

A composite biomarker score was then constructed to integrate evidence from expression trends, sex-dimorphic effects, disease-state differential expression, panel membership, network hub status, Open Targets disease scores, eQTL support, fine-mapping, druggability and colocalisation. Genes were classified as actionable when they combined at least three evidence layers, an approved drug and GWAS colocalisation support with H4 *>* 0.8.

### 2.7 Sex-Specific Evidence Scoring for Actionable Biomarkers

To identify which actionable biomarkers were most strongly supported by sex-specific evidence, we developed a separate score on a 0–100 scale. This score was intentionally distinct from the global biomarker score and focused on six dimensions: the number of male- or female-specific Kruskal–Wallis tissues, the presence and strength of DESeq2 sex-by-age interactions, the breadth of sex main effects, concordance between statistical approaches, eQTL tissue specificity and druggability with colocalisation support. The purpose of this score was not to replace the global biomarker ranking, but to prioritize candidates for future sex-stratified experimental validation.

## 3 Results

### 3.1 Sex-Dimorphic Age Trends and Disease-State Concordance

Across 38,694 gene–tissue Kruskal–Wallis tests, 4,404 showed significant age-associated expression trends at BH-FDR *<* 0.05 (11.4%), and 1,305 remained significant after Bonferroni correction (3.4%). Sex-specific classification revealed a strong male bias, with 2,718 male-specific trends, 202 female-specific trends and 742 shared trends. This imbalance is partly attributable to the male-enriched GTEx sample composition, which provides greater statistical power in male-stratified analyses. However, the size of the difference also supports the presence of substantial biological sex dimorphism in age-related CVD gene expression (Figure 1). Among tissue-specific findings, *GDF15* increased significantly with age in male left ventricular tissue (padj = 0.0045), consistent with its role as a cardiac aging biomarker [30, 31], whereas *PRDX2* decreased in the same tissue, suggesting a possible decline in oxidative stress defence with cardiac aging.

**Figure 1:**
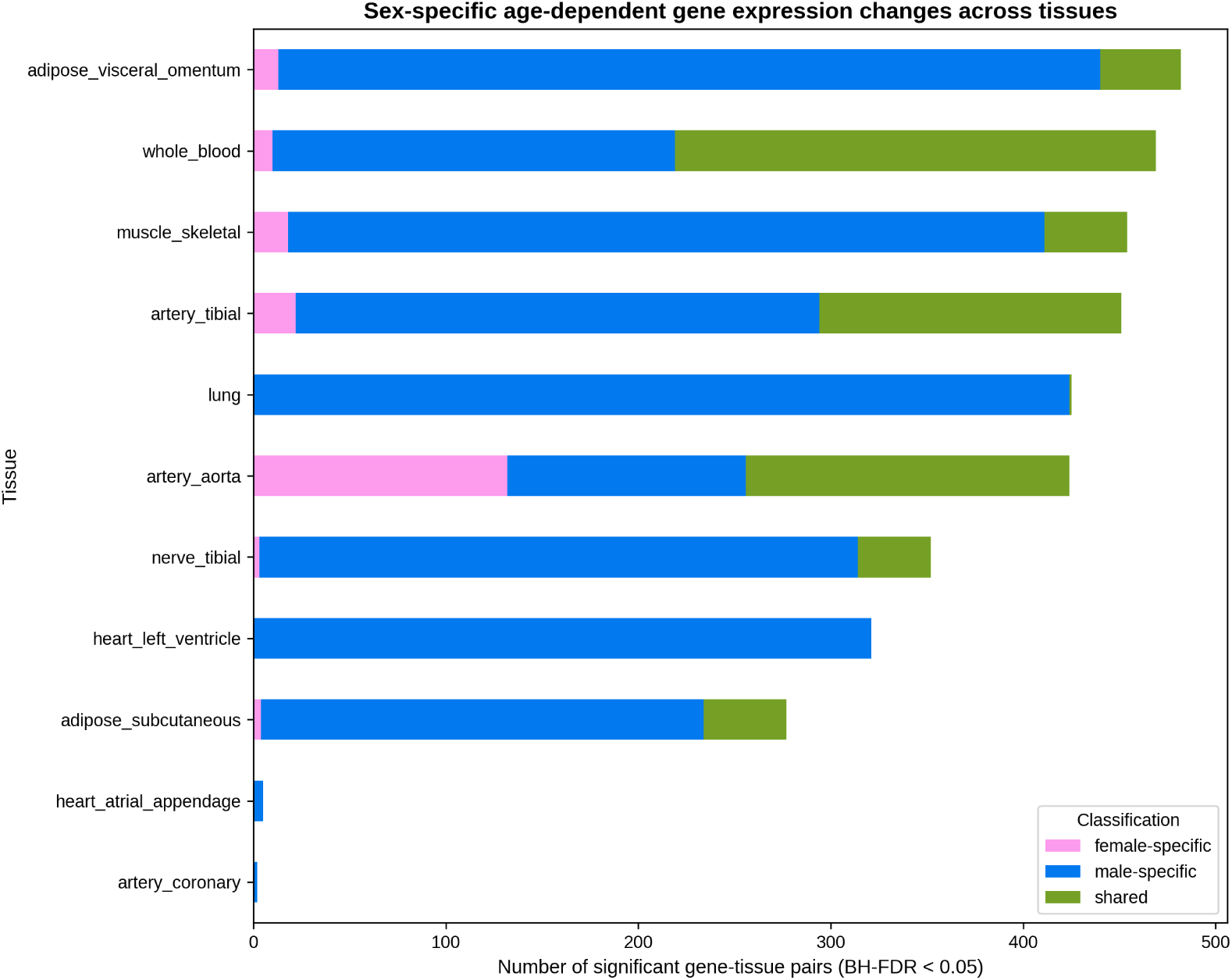
Summary of Kruskal–Wallis results across 17 tissues. Left: significant gene–tissue tests by correction method. Right: classification of significant trends by sex specificity.

DESeq2 interaction modelling identified 2,971 significant age main effects, 2,644 sex main effects and 87 significant sex-by-age interactions (padj *<* 0.05). Prominent interaction genes included *REN*, *MYL2*, *SELE*, *APOE*, *IL6* and *IL1B* (Figure 2). The strongest interaction involved *REN* (padj = 1.1 × 10*^−^*^6^), which showed a female-specific age effect in adipose tissue. This finding is consistent with the known sex dependence of renin–angiotensin–aldosterone system biology [32, 33, 34]. *APOE* also showed sex-dimorphic aging in tibial artery, in line with evidence that APOE-related vascular risk is modified by sex [35, 36].

**Figure 2:**
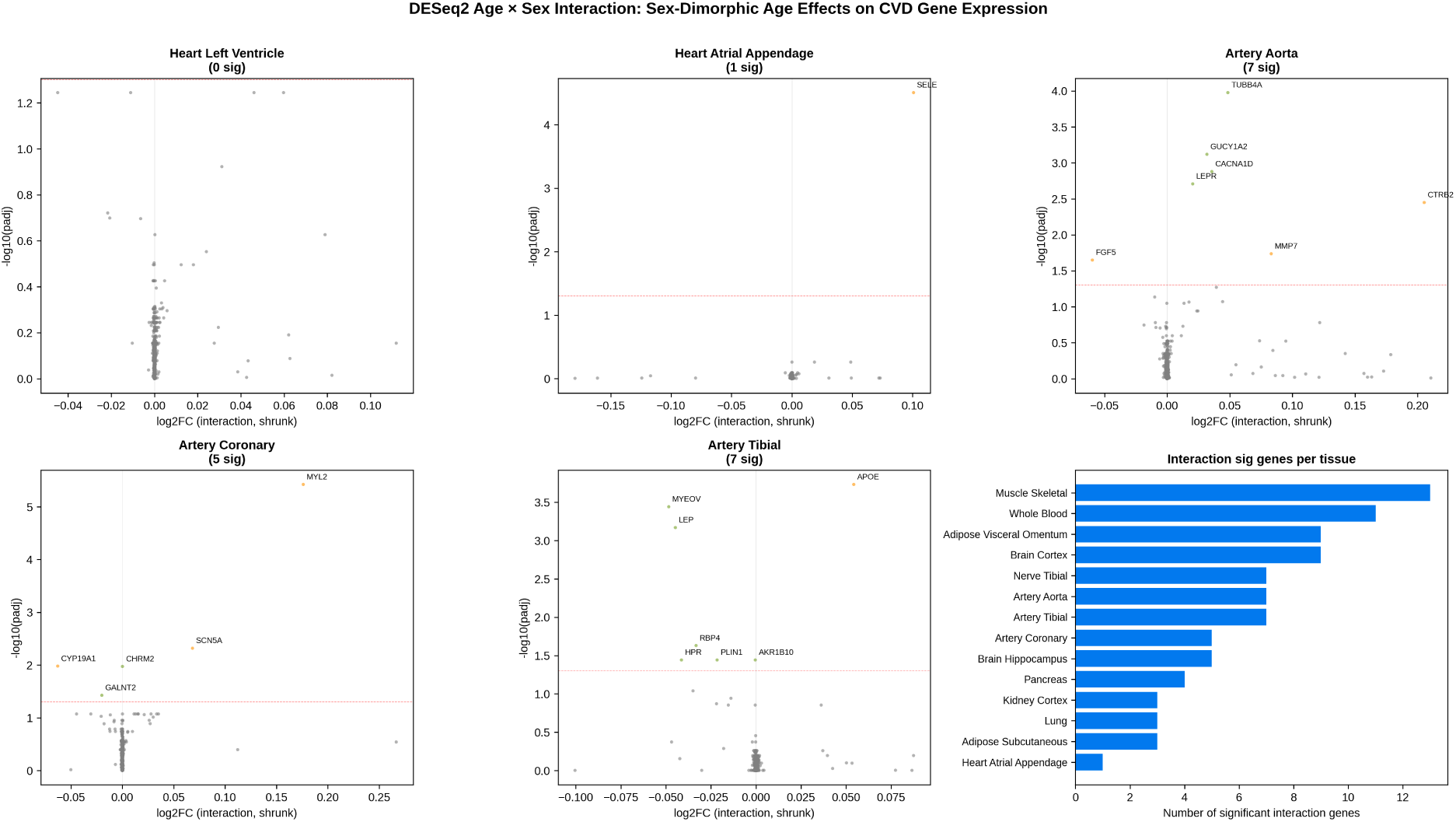
DESeq2 sex-by-age interaction results. The *x*-axis shows the shrunken log_2_ fold change of the interaction coefficient, and the *y*-axis shows the − log_10_ adjusted *p*-value. Significant genes are highlighted.

Combining the non-parametric and interaction-based analyses yielded 35 high-confidence sex-dimorphic genes, including 25 male-specific, 7 female-specific and 3 shared genes. Key concordant candidates included *REN*, *APOE*, *SRD5A2*, *GUCY1A2*, *LEPR*, *PGR* and *TF*. Of particular interest, *GUCY1A2*, a component of soluble guanylate cyclase involved in nitric oxide signaling, showed female-specific aging concordance and aligned with cGMP–PKG pathway enrichment observed in the network analyses (Figure 3).

**Figure 3:**
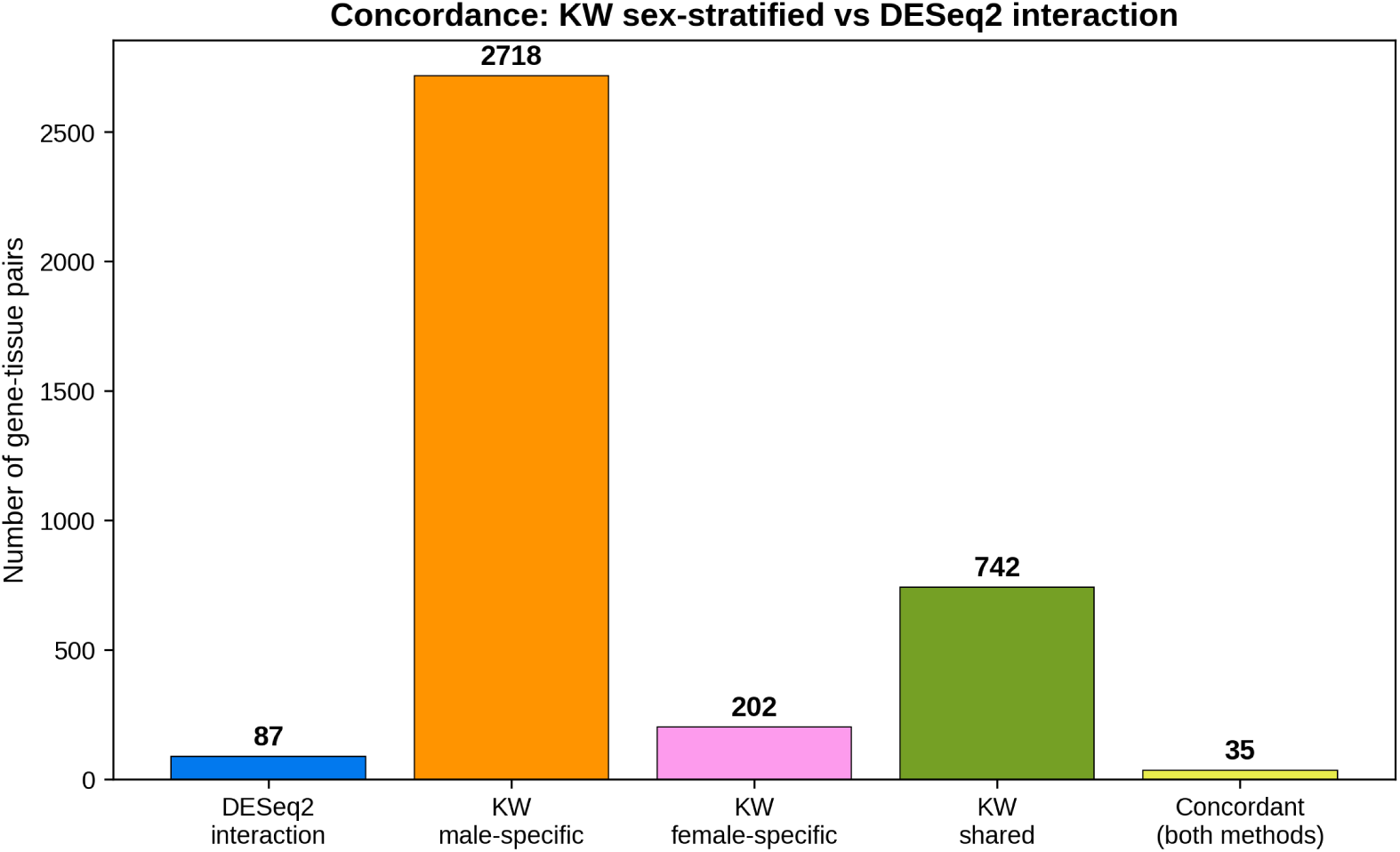
Concordance between Kruskal–Wallis and DESeq2 interaction results, showing the overlap of sex-dimorphic genes identified by the two approaches.

Disease-state validation further supported the relevance of the basal GTEx signatures. GSE116250 identified 2,191 differentially expressed genes in failing versus non-failing left ventricular samples, whereas GSE55296 identified 5 differentially expressed genes in ischemic or dilated cardiomyopathy, a more limited result likely reflecting smaller sample size. Intersecting disease-state genes with the CVD candidate set identified 118 genes that were both age- or sex-trended in GTEx and dysregulated in disease. Fourteen sex-dimorphic genes were also differentially expressed in disease, including *GDF15*, *ACACB*, *CXCR4*, *LPL* and *RBP4* (Figure 4).

**Figure 4:**
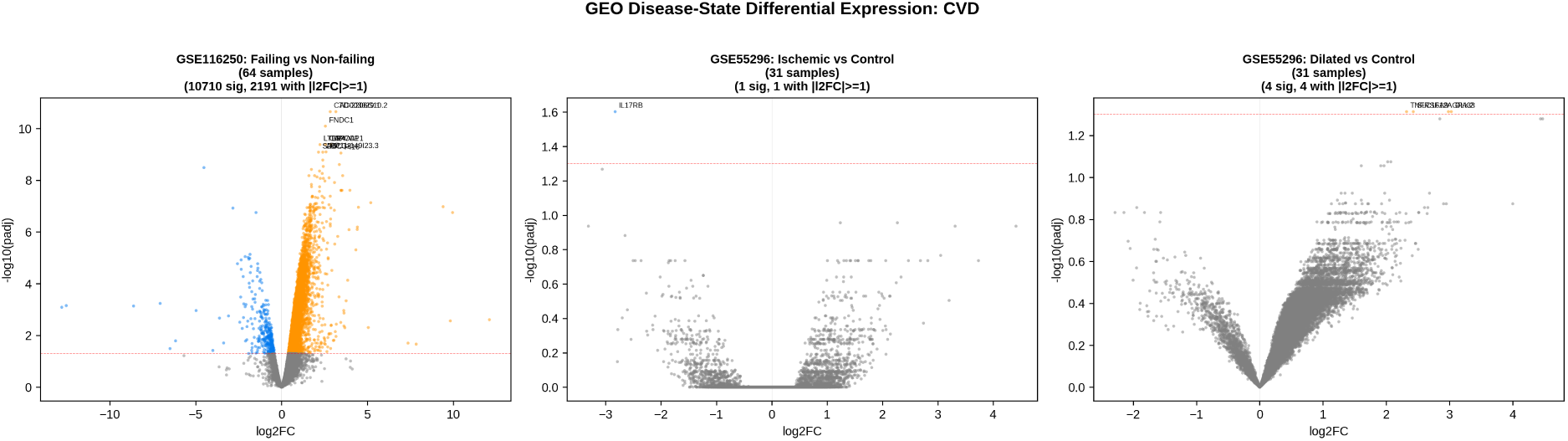
Disease-state differential expression analysis in GEO heart datasets. Left: failing versus non-failing heart samples. Center/right: ischemic or dilated cardiomyopathy versus control samples.

### 3.2 Network Topology, Ricci Curvature, and Sex-Stratified Geometry

The BioGRID and STRING CVD networks had comparable sizes but different organizational properties. BioGRID was denser and less fragmented, whereas STRING displayed substantially higher clustering. BioGRID was disassortative, indicating that hubs tend to connect with lower-degree nodes, while STRING was assortative, consistent with more locally clustered pathway structure [22]. Both networks showed strong small-world organization [24], had the same diameter and contained well-defined communities (Figure 5, Table 1). These differences are consistent with the complementary nature of the two resources: BioGRID emphasizes curated physical interactions, whereas STRING integrates multiple sources of functional evidence.

**Figure 5:**
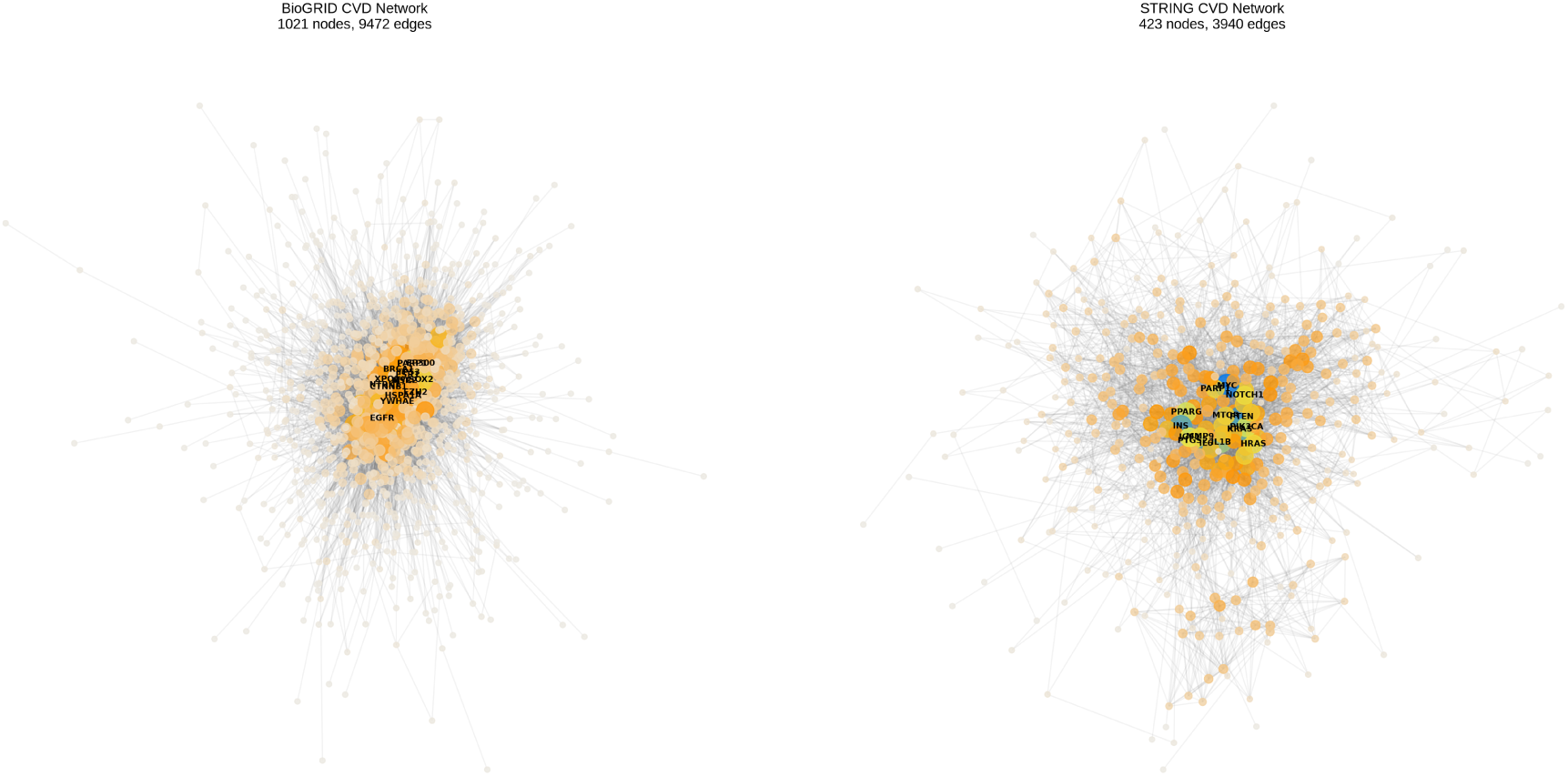
Network visualization comparison. Left: BioGRID CVD network. Right: STRING CVD network. Node size and color reflect degree; top hubs are labeled.

**Table 1:** Network geometry metrics for BioGRID and STRING CVD networks (largest connected component). *δ* = Gromov *δ*-hyperbolicity; *κ_G_* = combinatorial Gaussian curvature; *σ* = small-world index.

| Network | Nodes | Edges | Assort. | Modularity | $\sigma$ | Diam. | $\delta$ mean | $\kappa_G$ mean | Rich-club |
| --- | --- | --- | --- | --- | --- | --- | --- | --- | --- |
| BioGRID | 1,021 | 9,472 | $-0.108$ | 0.303 | 11.31 | 7 | 0.199 | 18.84 | 0.563 |
| STRING | 423 | 3,940 | $+0.107$ | 0.389 | 8.03 | 7 | 0.208 | 33.50 | 0.704 |

The most connected BioGRID nodes included *MYC*, *TP53*, *ESR1*, *EGFR* and *ESR2*, whereas the top STRING hubs included *AKT1*, *MYC*, *EGFR*, *CTNNB1* and *PTEN*. The centrality of *ESR1* and *ESR2* in BioGRID is noteworthy, because estrogen signaling is directly relevant to sex differences in vascular biology and cardiovascular aging.

Ricci curvature analysis revealed a clear geometric distinction between the networks. BioGRID had more negative Ollivier–Ricci curvature than STRING (mean = −0.053 vs. +0.026, Mann–Whitney *U*, *p <* 10*^−^*^260^), suggesting a greater abundance of bridge-like, hyperbolic edges. Forman–Ricci curvature showed the same pattern, with BioGRID displaying more negative values than STRING. In contrast, node-level combinatorial Gaussian curvature was positive in both networks and higher in STRING, reflecting its greater triangle density and local clustering.

Curvature was strongly linked to network function. Edges with high betweenness centrality, which connect otherwise distinct communities, had the most negative Ricci curvature. The correlation between Ollivier–Ricci curvature and edge betweenness was *ρ* = −0.566 in BioGRID and *ρ* = −0.697 in STRING. At the node level, Gaussian curvature correlated positively with degree and betweenness, indicating that hubs occupy positively curved regions, whereas bridge edges define negatively curved paths between communities (Figure 6).

**Figure 6:**
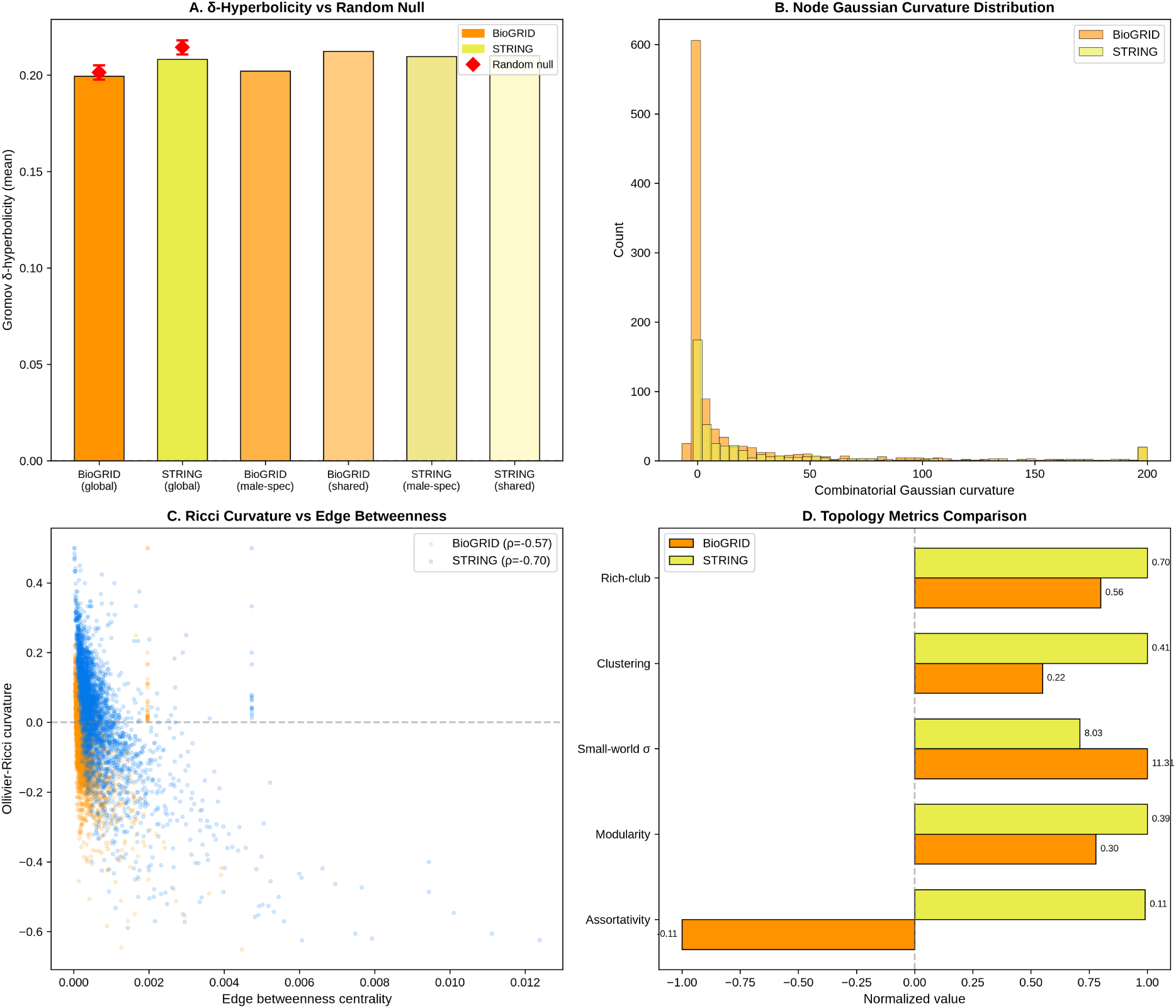
Network geometry metrics. (A) Gromov *δ*-hyperbolicity versus random null networks. (B) Combinatorial Gaussian curvature distribution. (C) Ollivier–Ricci curvature versus edge betweenness centrality. (D) Topology metric comparison.

Sex-stratified curvature analysis showed that shared sex-dimorphic genes occupied the most negatively curved positions in the BioGRID network (mean Ollivier–Ricci curvature = −0.098), suggesting that genes significant in both sexes may act as connectors between sex-biased functional modules. Shared-gene subnetworks also showed higher modularity than male-specific subnetworks, indicating a more cohesive community organization. Female-specific subnetworks were small, limiting geometric interpretation, but the available results support the idea that sex-dimorphic genes differ not only in expression pattern but also in network position (Table 2, Figures 7–8).

**Figure 7:**
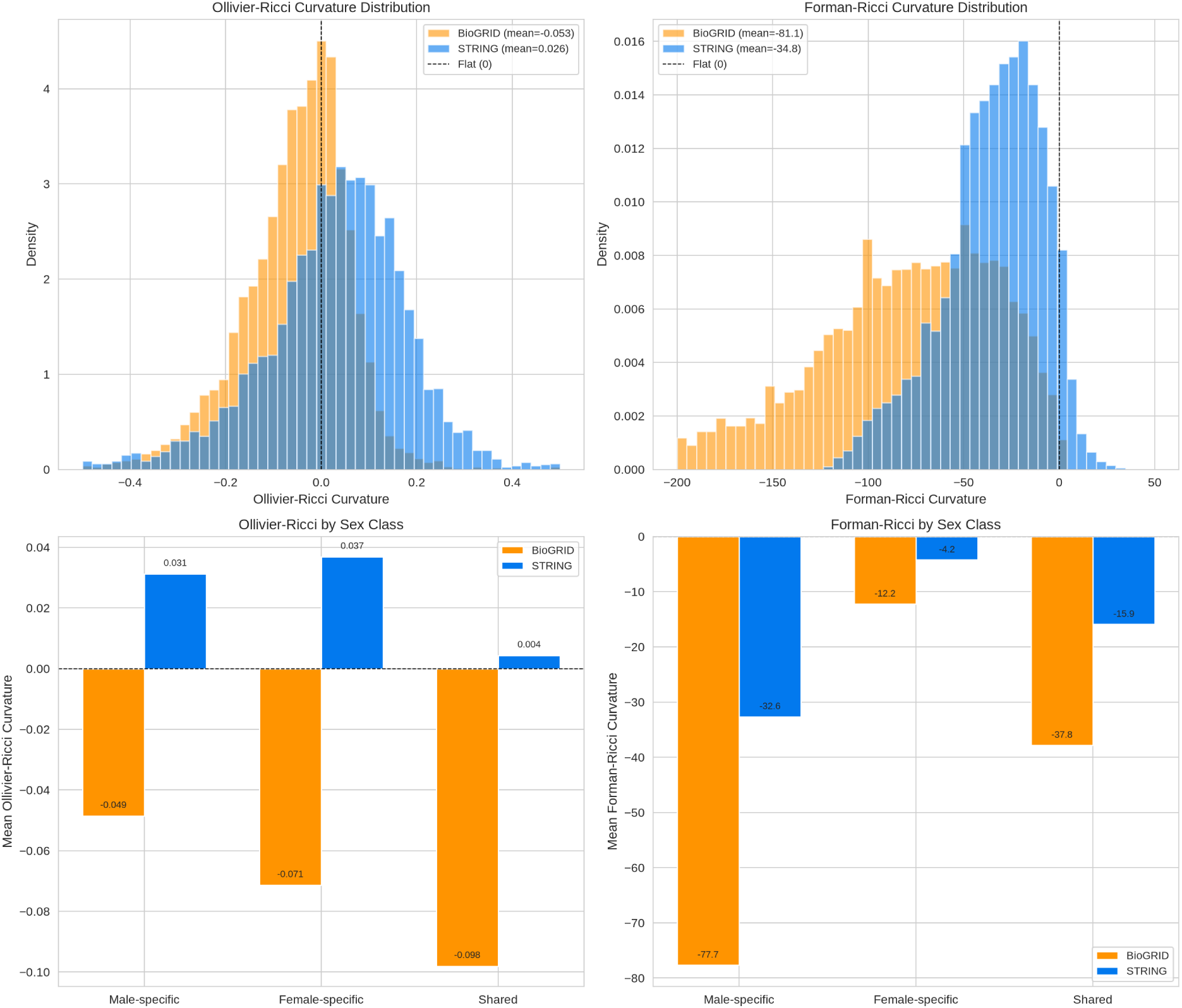
Ricci curvature comparison between BioGRID and STRING networks. Upper panels show distributions of Ollivier–Ricci and Forman–Ricci curvature. Lower panels summarize mean curvature by sex class.

**Figure 8:**
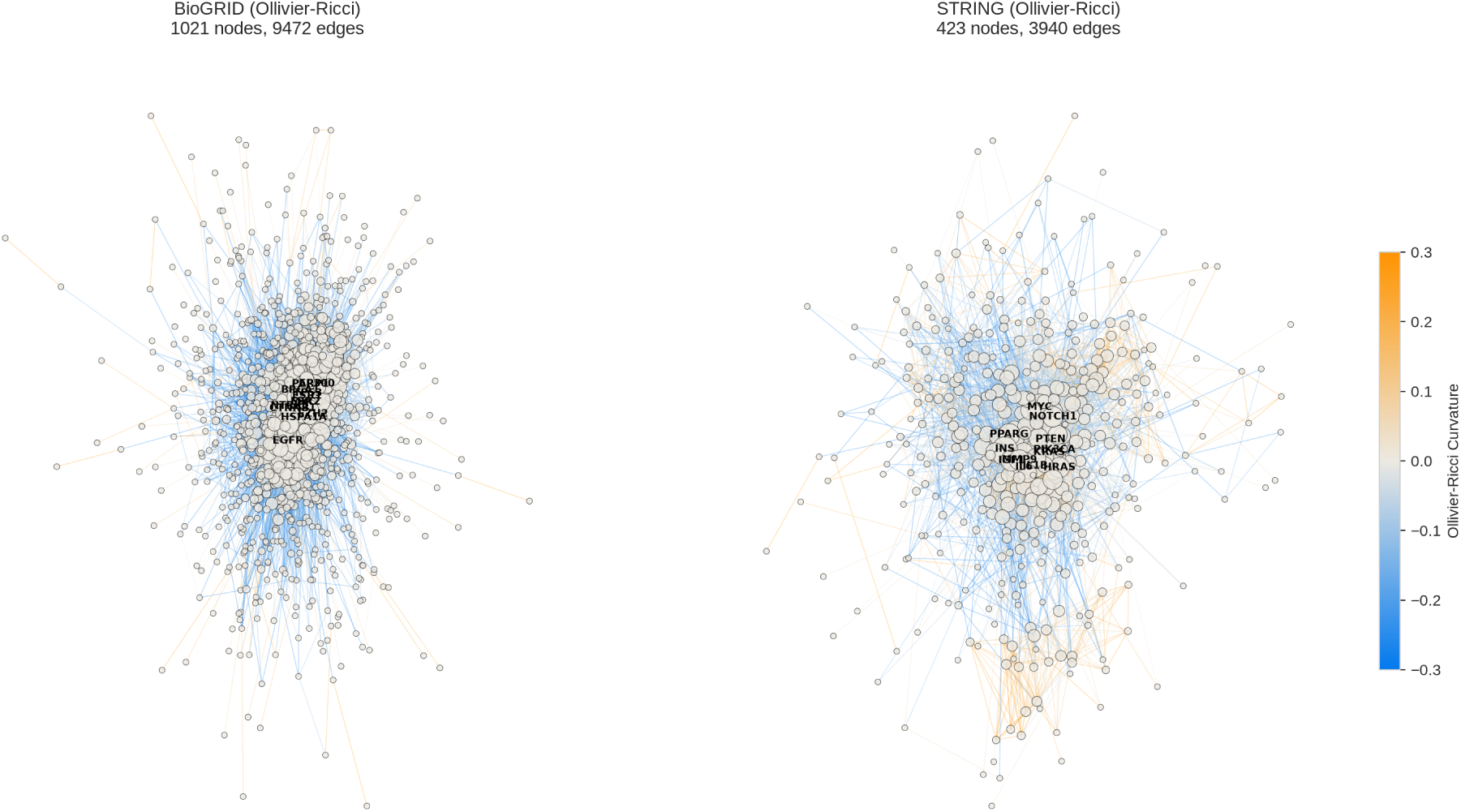
Network visualization with edges colored by Ollivier–Ricci curvature. Negative curvature identifies bridge-like edges, whereas positive curvature identifies locally clustered edges.

**Table 2:** Sex-stratified curvature and topology for BioGRID and STRING subnetworks.

| Network | Sex class | Nodes | Edges | Ollivier mean | Forman mean | Modularity | $\kappa_G$ mean |
| --- | --- | --- | --- | --- | --- | --- | --- |
| BioGRID | Male-specific | 757 | 5,797 | $-0.049$ | $-77.7$ | 0.319 | 11.66 |
| BioGRID | Shared | 130 | 311 | $-0.098$ | $-37.8$ | 0.465 | $-0.16$ |
| STRING | Male-specific | 312 | 2,370 | $+0.031$ | $-32.6$ | 0.382 | 22.07 |
| STRING | Shared | 64 | 132 | $+0.004$ | $-15.9$ | 0.478 | 0.14 |

### 3.3 Hyperbolic Geometry: Gromov *δ*-Hyperbolicity and Poincaré Disk Embedding

We next asked whether CVD interaction networks display hyperbolic organization, as expected for hierarchical biological networks [14]. Gromov *δ*-hyperbolicity showed that both networks were only modestly tree-like at the global level. BioGRID had *δ*_mean_ = 0.199 and STRING had *δ*_mean_ = 0.208, with both values close to degree-preserving random null models. The negative null-model *z*-scores suggested a slight tendency toward hyperbolicity, but not a strongly tree-like global structure.

In contrast, Poincaré disk embedding captured the graph structure substantially better than Euclidean embedding. Hyperbolic distances correlated strongly with shortest-path distances in both BioGRID (*ρ* = 0.712) and STRING (*ρ* = 0.692), whereas Euclidean spectral embedding performed poorly (*ρ* = −0.025 and 0.196, respectively). This indicates that, although the networks are not globally tree-like, their hierarchy, degree heterogeneity and functional community structure are naturally represented in hyperbolic space [15, 27].

Sex-stratified embeddings produced similarly high correlations, including *ρ* = 0.719 for male-specific BioGRID, *ρ* = 0.713 for shared BioGRID, *ρ* = 0.684 for male-specific STRING and *ρ* = 0.775 for the shared STRING subnetwork. These results suggest that hyperbolic organization is a stable property of CVD interaction networks and is preserved across sex-dimorphic gene subsets (Figure 9).

**Figure 9:**
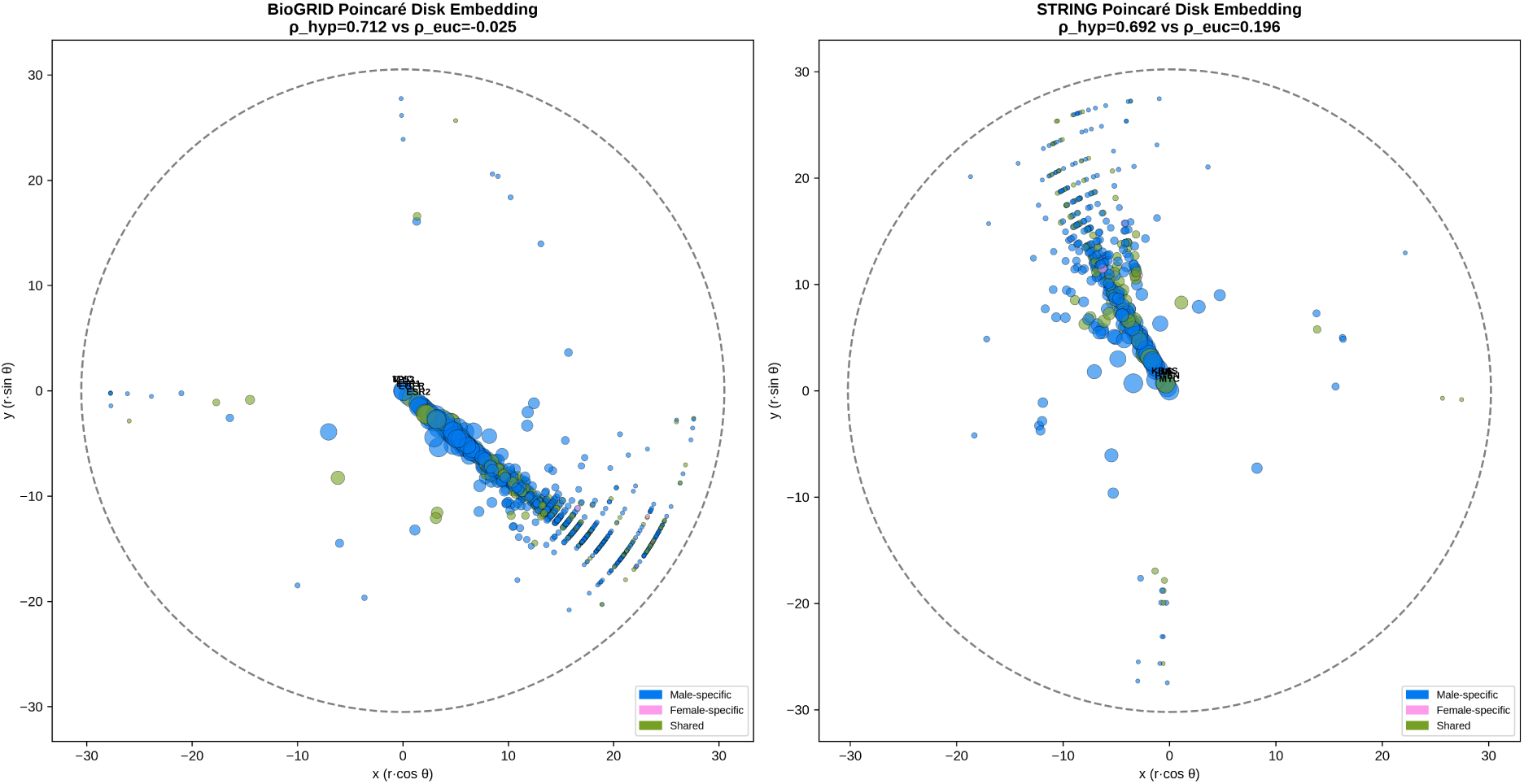
Poincaré disk embedding of CVD networks. Left: BioGRID. Right: STRING. Node colors indicate sex classification, and node size reflects degree. Top hub genes are labeled.

### 3.4 eQTL Regulatory Landscape: Effect Sizes, Tissue Specificity, and Fine-Mapping

cis-eQTL analysis showed that regulatory variation is widespread among CVD genes. Of the 1,176 candidate genes, 1,131 (96.2%) had at least one significant cis-eQTL in at least one selected tissue, corresponding to 8,710 gene–tissue associations. eQTL coverage varied by tissue and was correlated with sample size, with tibial nerve showing the largest number of CVD eGenes and kidney cortex the fewest.

Effect-size analysis revealed extensive tissue dependence. Across genes with eQTLs in at least two tissues, the mean tissue-specificity index was *τ* = 0.434, with 370 genes showing high tissue specificity and 82 showing broadly shared effects. Directional concordance was low: only 214 of 1,073 genes (19.9%) had eQTL effects with the same sign across all tissues. Thus, the same regulatory locus may increase expression in one tissue but decrease it in another, emphasizing the importance of tissue context when interpreting genetic regulation. The most tissue-specific genes included *ADGRG6*, *HSPA4* and *MEN1*, whereas *PSIP1*, *ADIPOQ* and *REN* showed more widespread regulatory patterns. Tissue clustering based on eQTL sharing placed most tissues together, with liver and kidney cortex emerging as outliers (Figure 12).

SuSiE fine-mapping identified 699 genes (59.4%) with high-PIP variants and 465 genes with very high-PIP variants. Nearly half of high-PIP associations belonged to single-variant credible sets, indicating strong resolution for causal variant prioritization. Cross-tissue analysis identified 352 variants that were high-PIP in at least two tissues, including variants recurrent across several cardiovascular and metabolic tissues. These variants represent robust regulatory signals that may be particularly informative for downstream functional validation.

Integrating regulatory and expression evidence showed strong convergence. Genes with eQTLs had a higher Kruskal–Wallis significance rate than genes without eQTLs, and the number of eQTL tissues correlated with the number of age-trend tissues. In contrast, eQTL breadth did not correlate with the number of DESeq2 sex-by-age interaction tissues, suggesting that sex-by-age effects may be driven by mechanisms beyond local germline regulation, such as hormonal, epigenetic or trans-regulatory processes. Six genes—*RBP4*, *TUBB4A*, *PDE5A*, *LEPR*, *SHROOM3* and *PTGIS* —showed evidence across all six analytical layers (Figures 11–12).

**Figure 10:**
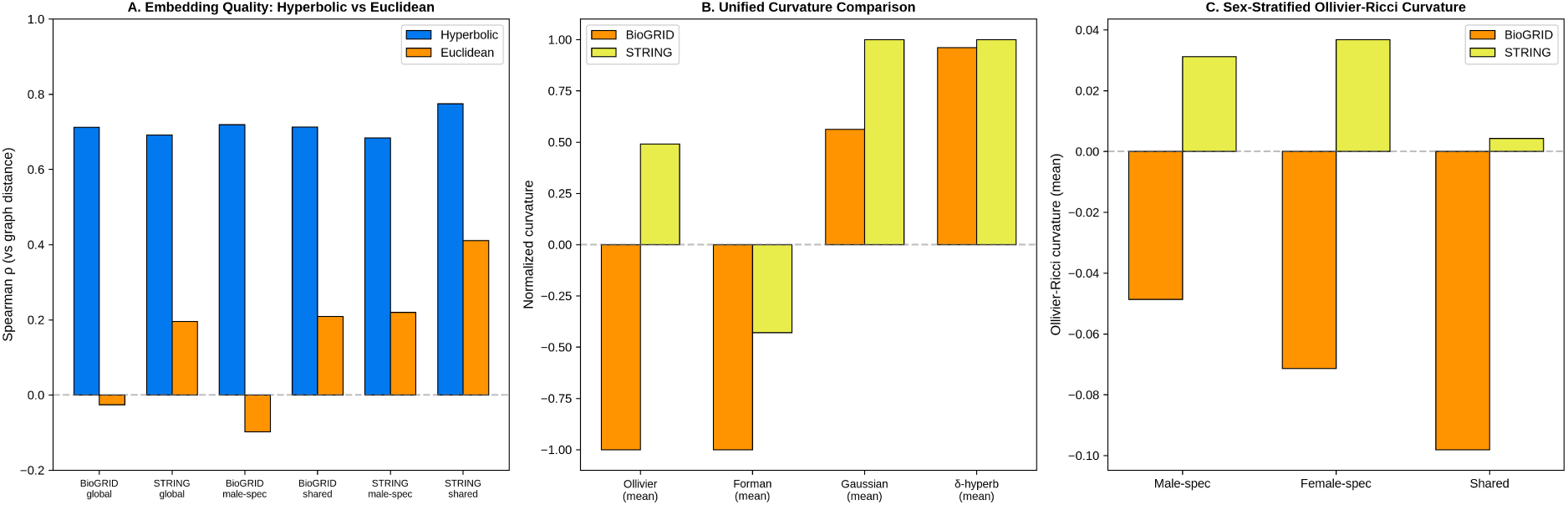
Hyperbolic geometry summary. (A) Embedding quality for hyperbolic and Euclidean spaces across global and sex-stratified subnetworks. (B) Curvature comparison across metrics. (C) Sex-stratified Ollivier–Ricci curvature.

**Figure 11:**
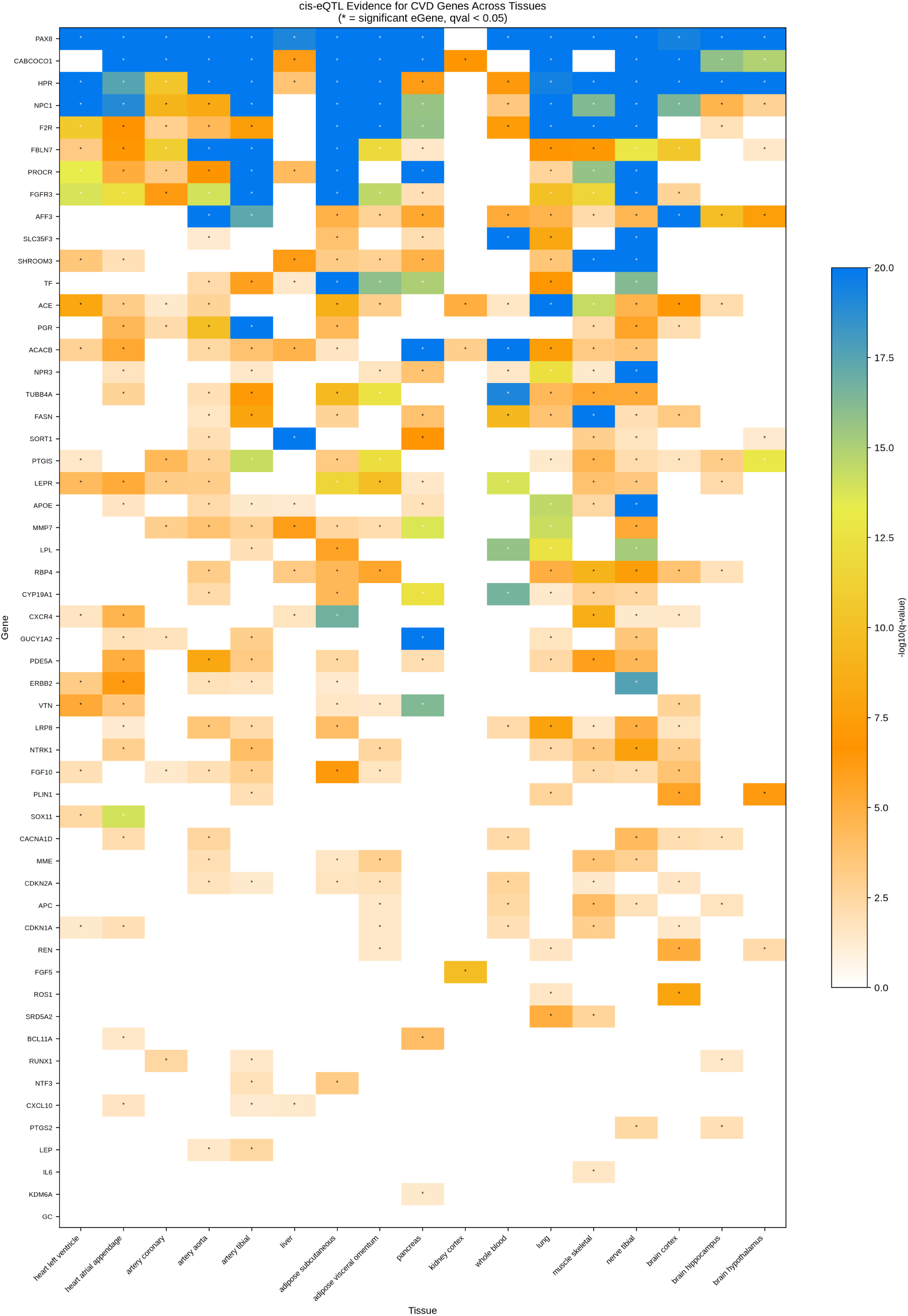
cis-eQTL evidence heatmap for top biomarker candidates and concordant sex-dimorphic genes across 17 tissues. Color intensity represents − log_10_(q-value); asterisks indicate significant eGenes.

### 3.5 Open Targets Druggability and GWAS Colocalisation

Open Targets integration provided therapeutic and genetic context for the CVD candidate set. Disease association scores were available for 910 genes in coronary artery disease and 827 genes in heart failure. In coronary artery disease, the highest-ranked targets included canonical genes such as *APOE*, *LDLR*, *GUCY1A1*, *PCSK9* and *HMGCR*, reflecting strong genetic and clinical evidence. In heart failure, top targets included genes related to neurohormonal signaling, consistent with established disease biology.

Drug and tractability analysis showed that 360 genes had at least one drug or clinical candidate, and 289 had approved drugs. The most heavily drugged targets included *DRD2*, *SLC6A2*, *ADRB2*, *HRH1* and tubulin-family genes. The recovery of established cardiovascular targets such as *PCSK9*, *ACE* and *HMGCR* supports the biological validity of the curated gene set and the prioritization strategy.

Genetic colocalisation further strengthened target interpretation. Among the top 50 biomarker candidates, 49 had credible-set evidence and 48 showed at least one GWAS colocalisation with H4 *>* 0.8. This indicates that, for most top-ranked genes, the same causal variant is likely to influence both molecular regulation and disease association. Genes with many colocalisation events included *SORT1*, *ERBB2*, *ACE*, *LPL* and *MME*, providing genetic support beyond expression association alone (Figure 13).

**Figure 12:**
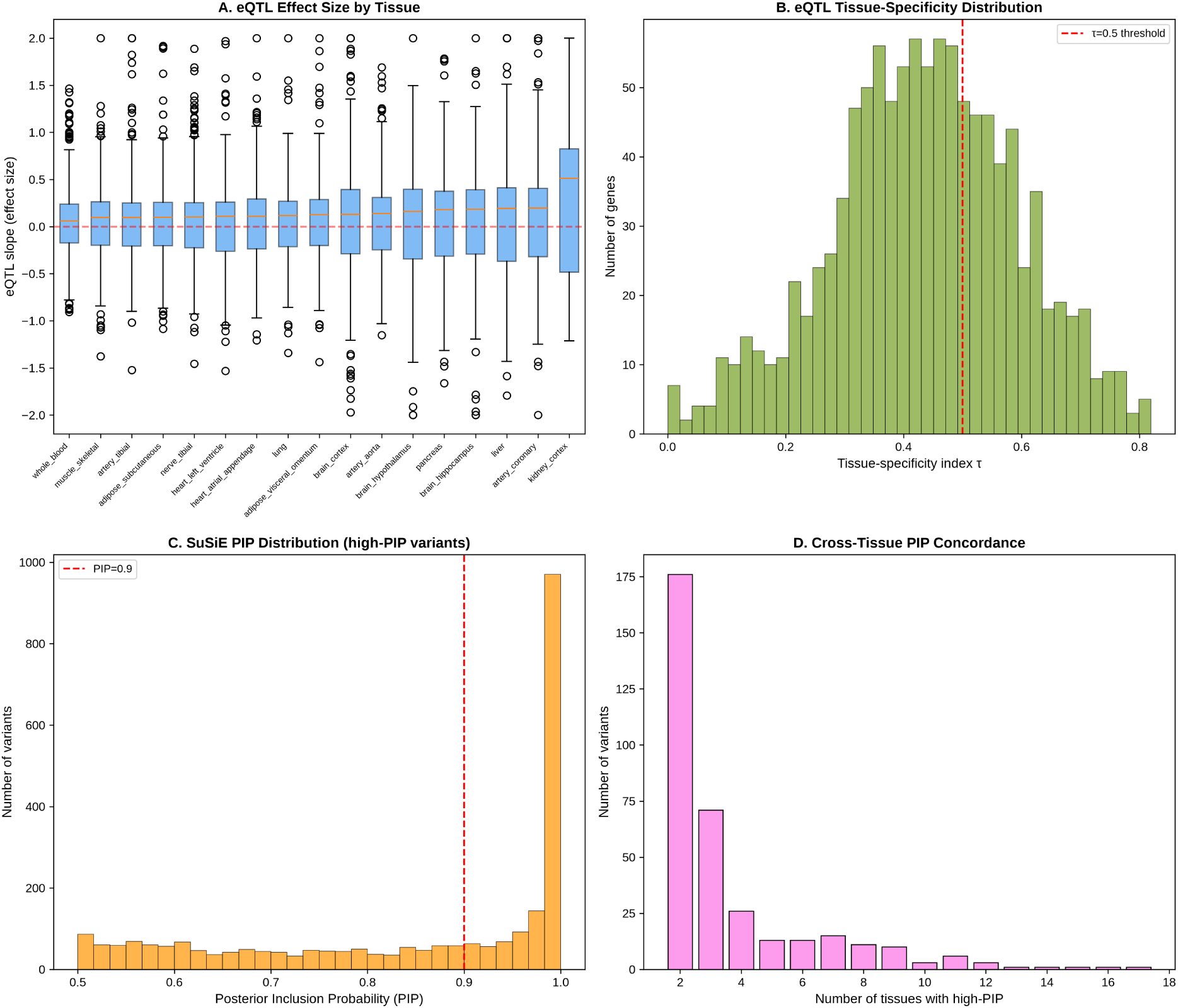
Extended eQTL analysis. (A) eQTL effect-size distribution by tissue. (B) Tissue-specificity index distribution. (C) SuSiE PIP distribution for high-PIP variants. (D) Cross-tissue recurrence of high-PIP variants.

**Figure 13:**
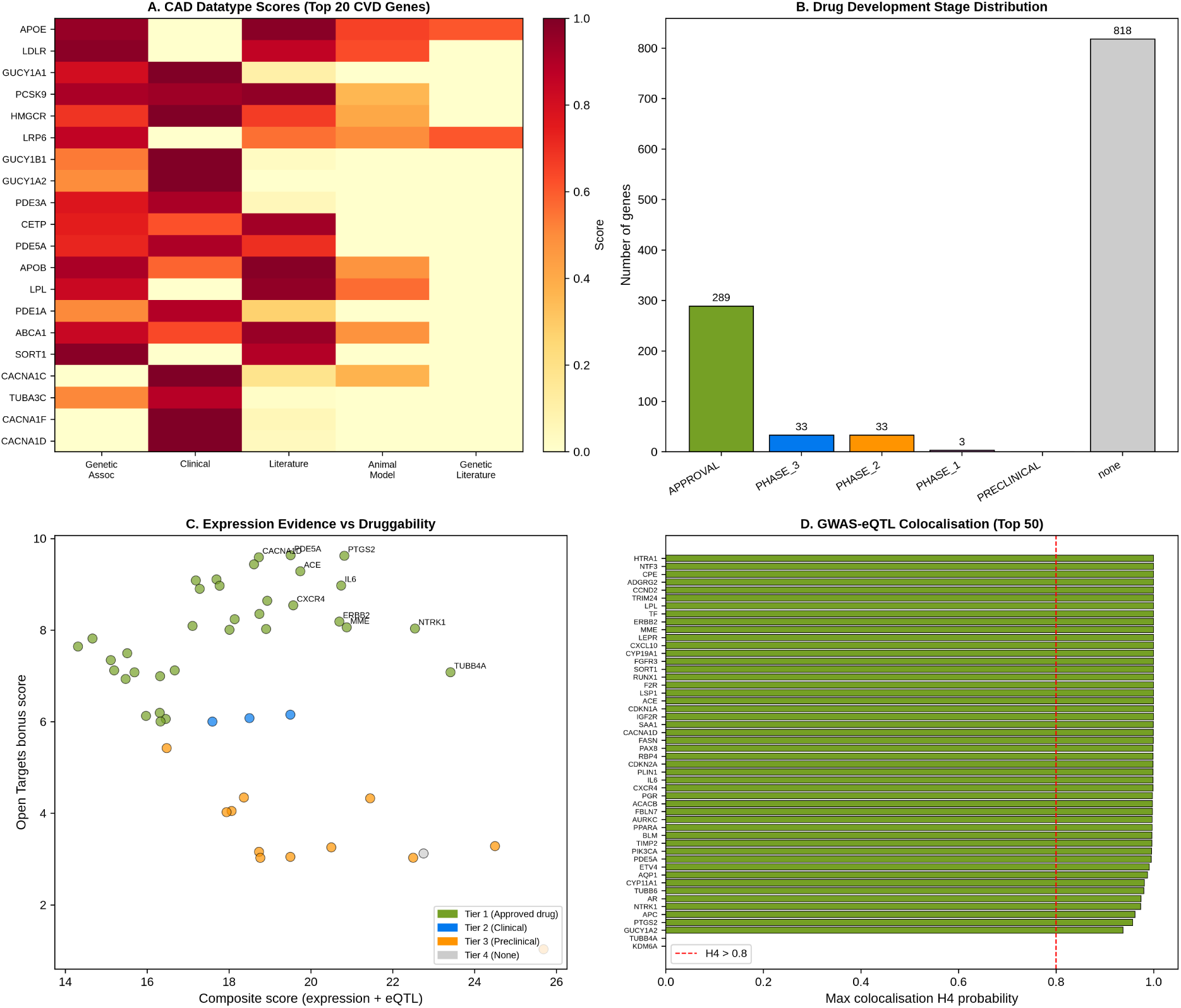
Open Targets integration. (A) Coronary artery disease evidence scores for top CVD genes. (B) Drug development stage distribution. (C) Relationship between molecular evidence and druggability. (D) GWAS–eQTL colocalisation probability for top biomarker candidates.

### 3.6 Integrated Biomarker Ranking and Actionable Targets

The final biomarker score integrated expression trends, sex-dimorphic evidence, disease-state differential expression, network centrality, eQTL support, SuSiE fine-mapping, Open Targets disease scores, druggability and GWAS colocalisation. The highest-ranking candidates were *NTRK1*, *TUBB4A*, *PTGS2*, *IL6*, *PDE5A*, *ACE*, *MME*, *ERBB2*, *CACNA1D* and *CXCR4* (Table 3, Figure 14).

**Figure 14:**
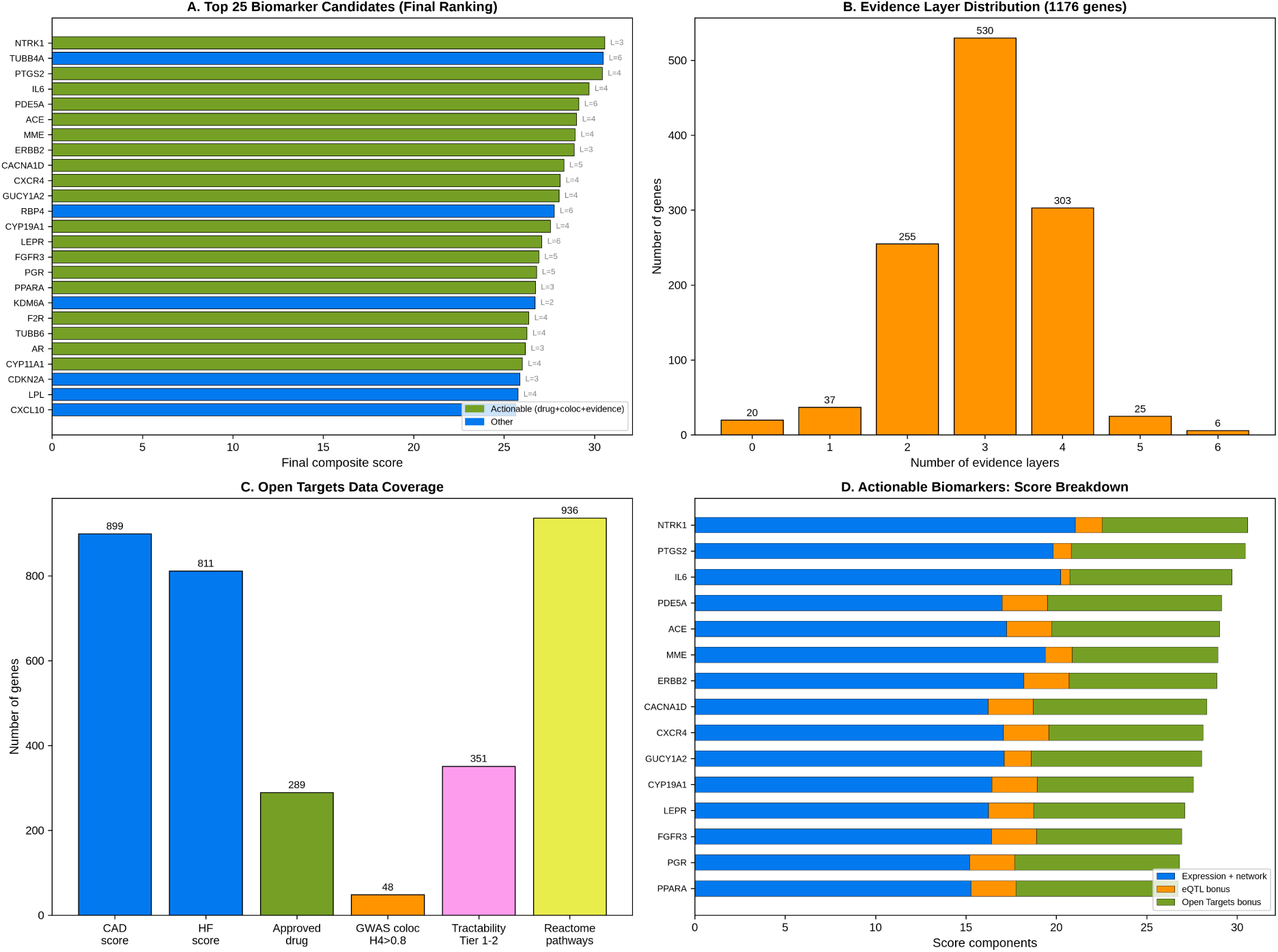
Final integrated biomarker ranking. (A) Top 25 candidates by final composite score.(B) Evidence layer distribution. (C) Open Targets data coverage. (D) Actionable biomarker score breakdown.

**Table 3:** Top 15 biomarker candidates by final composite score, integrating expression, eQTL, network, and Open Targets evidence. Actionable = ≥ 3 evidence layers + approved drug + GWAS coloc H4 *>* 0.8.

| Rank | Gene | Score | Layers | CAD score | Tract. tier | Drugs | Drug stage | Coloc H4 |
| --- | --- | --- | --- | --- | --- | --- | --- | --- |
| 1 | *NTRK1* | 30.58 | 3 | 0.005 | 1 | 17 | APPROVAL | 0.974 |
| 2 | *TUBB4A* | 30.49 | 6 | 0.538 | 1 | 88 | APPROVAL | 0.000 |
| 3 | *PTGS2* | 30.44 | 4 | 0.598 | 1 | 79 | APPROVAL | 0.957 |
| 4 | *IL6* | 29.71 | 4 | 0.414 | 1 | 6 | APPROVAL | 0.999 |
| 5 | *PDE5A* | 29.14 | 6 | 0.669 | 1 | 13 | APPROVAL | 0.996 |
| 6 | *ACE* | 29.02 | 4 | 0.603 | 1 | 24 | APPROVAL | 1.000 |
| 7 | *MME* | 28.93 | 4 | 0.027 | 1 | 4 | APPROVAL | 1.000 |
| 8 | *ERBB2* | 28.88 | 3 | 0.079 | 1 | 48 | APPROVAL | 1.000 |
| 9 | *CACNA1D* | 28.32 | 5 | 0.607 | 1 | 41 | APPROVAL | 0.999 |
| 10 | *CXCR4* | 28.11 | 4 | 0.259 | 1 | 8 | APPROVAL | 0.999 |
| 11 | *GUCY1A2* | 28.05 | 4 | 0.680 | 1 | 15 | APPROVAL | 0.938 |
| 12 | *RBP4* | 27.78 | 6 | 0.115 | 3 | 0 | none | 0.999 |
| 13 | *CYP19A1* | 27.57 | 4 | 0.302 | 1 | 6 | APPROVAL | 1.000 |
| 14 | *LEPR* | 27.09 | 6 | 0.127 | 1 | 2 | APPROVAL | 1.000 |
| 15 | *FGFR3* | 26.93 | 5 | 0.005 | 1 | 25 | APPROVAL | 1.000 |

Nineteen genes met the criteria for actionable biomarkers, defined by convergence across at least three evidence layers, the presence of an approved drug and GWAS colocalisation support. This group included *NTRK1*, *PTGS2*, *IL6*, *PDE5A*, *ACE*, *MME*, *ERBB2*, *CACNA1D*, *CXCR4*, *GUCY1A2*, *CYP19A1*, *LEPR*, *FGFR3*, *PGR*, *PPARA*, *F2R*, *TUBB6*, *AR* and *CYP11A1*. Among these, *PDE5A* and *LEPR* were supported by all six evidence layers, making them especially strong candidates at the intersection of expression, regulatory, disease and therapeutic evidence.

The integrated score also changed the prioritization relative to expression-only evidence. Genes with strong expression signals but limited therapeutic or genetic support were ranked lower, whereas druggable and genetically supported genes moved upward. For example, *NTRK1*, *PTGS2* and *ACE* rose in rank because their molecular evidence converged with approved drugs and colocalisation support. This shift illustrates the value of integrating biological discovery with translational evidence when prioritizing candidate biomarkers.

### 3.7 Sex-Specific Evidence Ranking of Actionable Biomarkers

Because the global biomarker score captures overall evidence convergence, we next asked which actionable genes were most specifically supported by sex-dimorphic evidence. The sex-specific evidence score separated the 19 actionable biomarkers into three tiers based on sex-stratified age trends, interaction effects, cross-method concordance, regulatory support and druggability (Table 4, Figure 15).

**Figure 15:**
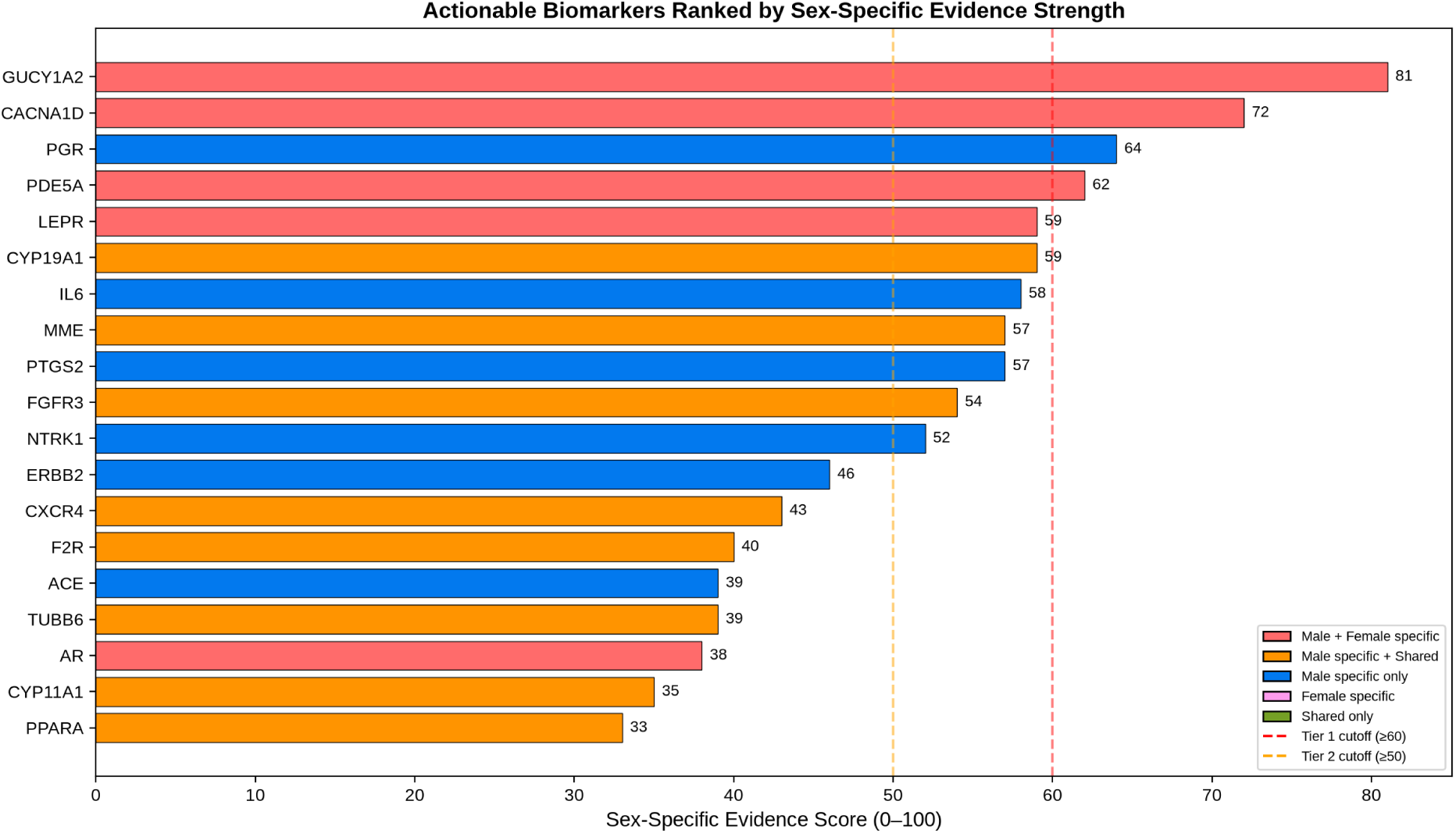
Sex-specific evidence ranking of the 19 actionable biomarkers. Bar length represents the composite sex-specific evidence score. Dashed lines mark Tier 1 and Tier 2 cutoffs.

**Table 4:** Sex-specific evidence ranking of the 19 actionable biomarkers. Score components: KW = sex-specific KW tissues (max 25), Int = DESeq2 interaction (max 25), SexMain = sex main effect breadth (max 15), Conc = cross-layer concordance (max 15), eQTL = eQTL tissue-specificity (max 10), Drug/Coloc = druggability and colocalisation (max 10). M = male-specific tissues, F = female-specific, S = shared.

| Tier | Gene | Total<br>/100 | M | F | S | Int sig<br>tissues | $\tau$ | KW<br>/25 | Int<br>/25 | SexMain<br>/15 | Conc<br>/15 | eQTL<br>/10 | Drug<br>/10 |
| --- | --- | --- | --- | --- | --- | --- | --- | --- | --- | --- | --- | --- | --- |
| 1 | *GUCY1A2* | 81 | 3 | 2 | 0 | 2 | 0.577 | 15 | 21 | 10 | 15 | 10 | 10 |
| 1 | *CACNA1D* | 72 | 4 | 1 | 1 | 2 | 0.644 | 15 | 16 | 6 | 15 | 10 | 10 |
| 1 | *PGR* | 64 | 6 | 0 | 0 | 1 | 0.424 | 18 | 8 | 8 | 15 | 5 | 10 |
| 1 | *PDE5A* | 62 | 3 | 1 | 1 | 1 | 0.399 | 12 | 8 | 12 | 15 | 5 | 10 |
| 1 | *LEPR* | 59 | 4 | 1 | 0 | 1 | 0.206 | 15 | 8 | 6 | 15 | 5 | 10 |
| 1 | *CYP19A1* | 59 | 3 | 0 | 2 | 1 | 0.500 | 9 | 8 | 12 | 10 | 10 | 10 |
| 1 | *IL6* | 58 | 5 | 0 | 0 | 1 | — | 15 | 8 | 14 | 10 | 1 | 10 |
| 1 | *MME* | 57 | 2 | 0 | 2 | 2 | 0.278 | 6 | 16 | 10 | 10 | 5 | 10 |
| 1 | *PTGS2* | 57 | 5 | 0 | 0 | 1 | 0.231 | 15 | 8 | 12 | 10 | 2 | 10 |
| 2 | *FGFR3* | 54 | 2 | 0 | 1 | 1 | 0.543 | 6 | 8 | 10 | 10 | 10 | 10 |
| 2 | *NTRK1* | 52 | 6 | 0 | 0 | 0 | 0.541 | 18 | 0 | 14 | 0 | 10 | 10 |
| 2 | *ERBB2* | 46 | 7 | 0 | 0 | 0 | 0.453 | 21 | 0 | 10 | 0 | 5 | 10 |
| 3 | *CXCR4* | 43 | 5 | 0 | 2 | 0 | 0.534 | 15 | 0 | 8 | 0 | 10 | 10 |
| 3 | *F2R* | 40 | 4 | 0 | 1 | 0 | 0.580 | 12 | 0 | 8 | 0 | 10 | 10 |
| 3 | *ACE* | 39 | 3 | 0 | 0 | 0 | 0.411 | 9 | 0 | 15 | 0 | 5 | 10 |
| 3 | *TUBB6* | 39 | 3 | 0 | 2 | 0 | 0.702 | 9 | 0 | 10 | 0 | 10 | 10 |
| 3 | *AR* | 38 | 3 | 1 | 1 | 0 | 0.699 | 12 | 0 | 6 | 0 | 10 | 10 |
| 3 | *CYP11A1* | 35 | 4 | 0 | 1 | 0 | 0.387 | 12 | 0 | 8 | 0 | 5 | 10 |
| 3 | *PPARA* | 33 | 2 | 0 | 5 | 0 | 0.231 | 6 | 0 | 12 | 0 | 5 | 10 |

Tier 1 contained nine genes with the strongest and most concordant sex-dimorphic profiles. *GUCY1A2* ranked first, supported by both male- and female-specific age trends, two significant sex-by-age interactions, tissue-specific eQTL regulation and GWAS colocalisation. *CACNA1D* showed a similar bidirectional pattern, with increasing age trends in male metabolic and vascular tissues and decreasing expression in female aorta. *PGR* displayed the broadest male-specific signature, including left ventricular tissue, whereas *PDE5A* and *LEPR* combined bidirectional sex specificity with strong therapeutic and genetic support.

A recurrent pattern among the top candidates was a female-specific age-related decrease in aortic tissue. *GUCY1A2*, *CACNA1D*, *PDE5A* and *LEPR* all showed decreasing expression with age in female aorta, while several showed increasing trends in male adipose or muscle tissues. This vessel-specific pattern suggests a convergent sex-dependent remodeling process that may be relevant to vascular aging, endothelial function and atherosclerotic risk.

Tier 2 genes, including *FGFR3*, *NTRK1* and *ERBB2*, had broad male-specific age-trend signals but lacked consistent interaction evidence. Tier 3 genes, including *CXCR4*, *F2R*, *ACE*, *TUBB6*, *AR*, *CYP11A1* and *PPARA*, had weaker sex-specific support despite in some cases being strong therapeutic targets. The contrast between *ACE*, which has excellent druggability and colocalisation but limited sex-by-age interaction evidence, and *GUCY1A2* or *PDE5A*, which combine actionable status with sex-specific expression, illustrates why a dedicated sex-evidence score is useful.

The Poincaré embedding provided additional biological interpretation. The top sex-evidence biomarkers tended to occupy peripheral positions in the BioGRID disk, whereas several top global biomarkers, including *NTRK1*, *AR* and *ERBB2*, were central hubs. This suggests that the strongest sex-specific candidates are not necessarily global network hubs, but rather context-dependent regulators whose effects may be tissue- and sex-specific. Most actionable biomarkers also clustered in a common angular sector, indicating functional coherence despite differences in network centrality (Figure 16).

**Figure 16:**
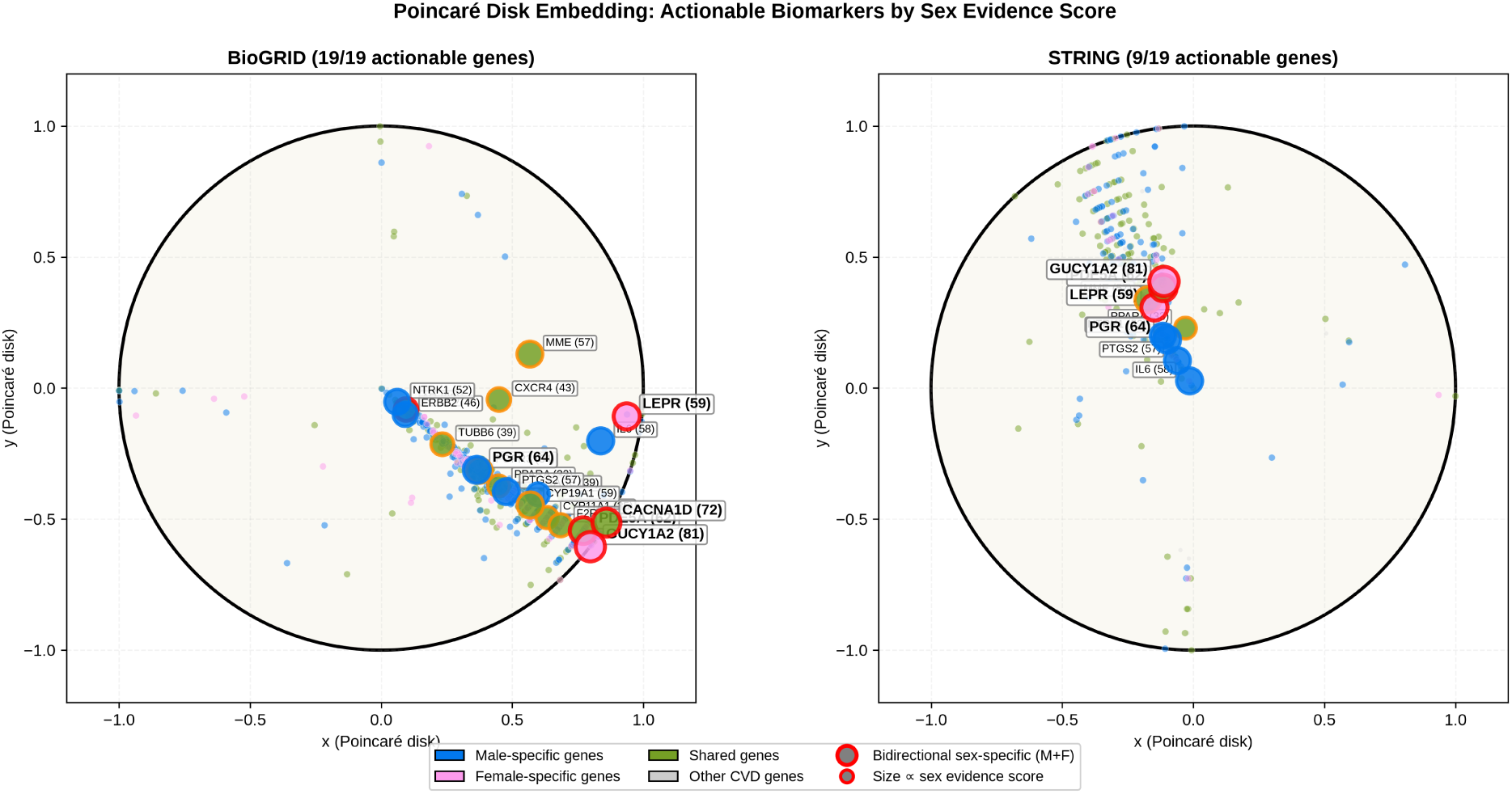
Poincaré disk embedding with actionable biomarkers highlighted. Node size is proportional to the sex-specific evidence score, and colors indicate sex classification. The separation between peripheral sex-specific biomarkers and central network hubs is visible.

## 4 Discussion

### 4.1 Sex-Dimorphic Aging, RAAS, and X-Chromosome Escapees

This study identifies widespread sex-dimorphic aging in the expression of CVD genes across cardiovascular, metabolic and neuroendocrine tissues. Although the male-biased GTEx sample composition likely contributes to the greater number of male-specific findings, the concordant set of 35 genes detected by both non-parametric and interaction-based approaches provides a robust basis for biological interpretation. Among these, *REN* was the strongest interaction signal, showing a female-specific age effect in adipose tissue. This result adds a tissue-specific dimension to the well-established sex dependence of the renin–angiotensin–aldosterone system, in which estrogen-related signaling and aging can shift the balance between vasodilatory and vasoconstrictive pathways [37, 33, 34].

These findings may have translational relevance. Sex-specific RAAS regulation is increasingly recognized as a contributor to differences in hypertension, heart failure, vascular aging and atherosclerotic disease between men and women. Although the present study was not designed to build clinical prediction models, the identification of sex-dependent age-related expression patterns suggests that molecular profiling could eventually refine cardiovascular risk stratification and therapeutic personalization.

The results also highlight the importance of genes whose sex effects are not explained solely by circulating hormones. *KDM6A*, an X-chromosome escape gene, ranked highly in expression-based analyses and is known to influence cardiometabolic biology. Its lower position in the final actionable ranking, due to limited druggability and colocalisation evidence, illustrates an important distinction: some genes may be mechanistically central to sex differences but not immediately actionable, whereas others, such as *ACE*, *PTGS2* and *PDE5A*, represent points where sex-dimorphic biology intersects with existing therapeutic opportunities.

More broadly, these results support the view that biological sex should be treated as a fundamental molecular variable in cardiovascular research. Future validation in balanced, prospective cohorts will be essential to determine whether these signatures can improve risk prediction, guide treatment selection or identify mechanisms underlying sex-specific disease phenotypes.

### 4.2 Network Geometry: From Ricci Curvature to Hyperbolic Embedding

The network analysis provides a geometric view of CVD biology. At the edge level, Ollivier–Ricci and Forman–Ricci curvature distinguished BioGRID from STRING, reflecting their different construction principles. BioGRID, which emphasizes curated physical interactions, contained more negatively curved bridge-like edges, whereas STRING, which integrates broader functional evidence, showed more clustered and positively curved structure. The strong negative correlation between Ricci curvature and edge betweenness confirms that negatively curved edges often serve as communication routes between communities [38, 39].

At the global level, Gromov hyperbolicity was close to random expectation, indicating that the networks are not simply tree-like. However, Poincaré embedding revealed that graph distances are much better preserved in hyperbolic than Euclidean space. This is biologically plausible: CVD interaction networks contain both hierarchical organization and cross-community connections, a structure that is not fully captured by a single global hyperbolicity statistic but is well represented by hyperbolic embedding [14, 15].

The embedding also offers interpretable biological coordinates. The radial coordinate separates central hubs from peripheral effectors: global biomarkers such as *NTRK1*, *AR* and *ERBB2* sit close to the disk center, whereas the strongest sex-specific biomarkers, including *GUCY1A2*, *CACNA1D*, *PDE5A* and *LEPR*, are peripheral. This suggests that sex-specific biomarkers may act as tissue- or context-dependent regulators rather than universal network hubs. The angular coordinate, by contrast, captures functional relatedness; the clustering of most actionable biomarkers in a common angular sector suggests pathway coherence despite large differences in node degree.

Sex-stratified geometry further showed that shared sex-dimorphic genes occupy relatively bridge-like positions and form modular subnetworks. This may explain why they are detected consistently across statistical approaches. By contrast, female-specific subnetworks were too small for reliable geometric analysis, reinforcing the need for balanced cohorts to better resolve female-specific cardiovascular molecular structure.

### 4.3 eQTL Regulatory Architecture: Tissue Specificity and Cross-Tissue Concordance

The eQTL analysis shows that most candidate CVD genes are under detectable cis-regulatory control, but that regulatory effects are highly tissue dependent. The low cross-tissue direction concordance is particularly important: a variant associated with increased expression in one tissue may be associated with decreased expression elsewhere. This observation cautions against interpreting eQTL or GWAS evidence without considering tissue context.

Fine-mapping added variant-level resolution, identifying many genes with high-PIP variants and a large subset of recurrent regulatory variants across tissues. The association between eQTL breadth and age-trend evidence suggests that local genetic regulation contributes to age-associated expression patterns. In contrast, the lack of association with sex-by-age interaction evidence indicates that sex-dimorphic aging may also involve mechanisms beyond cis-regulation, including hormonal signaling, chromatin remodeling, immune context or trans-regulatory effects.

### 4.4 Sex-Specific Evidence Ranking: From Biomarker Candidates to Prioritized Validation Targets

The sex-specific evidence score identified *GUCY1A2*, *CACNA1D*, *PGR*, *PDE5A* and *LEPR* as the strongest actionable candidates for sex-stratified validation. These genes combine sex-specific expression evidence with regulatory, genetic and therapeutic support, making them more compelling for targeted follow-up than genes ranked highly only by global biomarker score.

The *GUCY1A2* –*PDE5A* pair is especially relevant because both genes participate in the nitric oxide–sGC–cGMP signaling axis. *GUCY1A2* encodes a soluble guanylate cyclase subunit involved in cGMP production, whereas *PDE5A* encodes the enzyme that degrades cGMP. Their paired sex-dimorphic aging pattern, including female-specific decreases in aortic tissue and male-biased increases in metabolic tissues, suggests that this pathway may be differentially regulated with age in men and women. This hypothesis is experimentally testable in sex-stratified vascular and smooth muscle models.

*CACNA1D* and *LEPR* broaden the candidate set beyond cGMP signaling. *CACNA1D* implicates calcium handling, a pathway central to vascular tone and cardiomyocyte function, while *LEPR* links sex-dimorphic cardiovascular aging to adipose and metabolic regulation. The contrast between these candidates and established targets such as *ACE*, which is highly actionable but less sex-specific in this analysis, underscores the usefulness of separating general therapeutic relevance from sex-specific molecular evidence.

### 4.5 From Biomarker Candidates to Actionable Therapeutic Targets

The integration of druggability and GWAS colocalisation shifts the analysis from biomarker discovery toward therapeutic prioritization. The 19 actionable biomarkers combine molecular evidence with approved drugs and genetic support, making them attractive candidates for sex-stratified cardiovascular investigation. Several have direct clinical relevance: *ACE* is targeted by ACE inhibitors, *PDE5A* by phosphodiesterase inhibitors used in pulmonary hypertension, *PTGS2* by COX-2 inhibitors with complex cardiovascular risk profiles, and *IL6* by anti-inflammatory strategies relevant to atherosclerosis [40].

The colocalisation evidence is particularly important because it supports a shared causal variant underlying both molecular regulation and disease association [17, 18]. Genes with strong colocalisation, such as *ACE*, *MME*, *ERBB2*, *LEPR* and *FGFR3*, are therefore stronger candidates for causal involvement than genes supported by expression association alone.

A strength of the present work is the integration of transcriptomic, regulatory, network and druggability evidence into a unified prioritization framework. Rather than identifying genes merely associated with CVD, this approach highlights candidates that are biologically plausible, genetically supported and potentially pharmacologically actionable. At the same time, the findings should be viewed as hypothesis-generating. The analyses are based on basal tissue transcriptomics and public datasets rather than prospective clinical cohorts, so the proposed biomarkers should not yet be interpreted as clinical diagnostic or prognostic tools. Their utility must be assessed in independent cohorts and experimental models.

Overall, the study provides a framework for prioritizing sex-dependent cardiovascular biomarkers and therapeutic targets. Established targets such as *PCSK9*, *HMGCR* and *ACE* provide internal validation, while candidates such as *GUCY1A2*, *PDE5A*, *CACNA1D* and *LEPR* nominate pathways for future sex-aware cardiovascular research.

### 4.6 Limitations

Several limitations should be acknowledged. First, GTEx v11 has an approximately 2:1 male-to-female sample ratio, which increases power for male-specific detections and limits interpretation of female-specific trends. Second, GTEx is based on bulk tissue RNA-seq; observed differences may therefore reflect changes in cell-type composition as well as within-cell expression. Third, the external disease datasets were limited in size and provided normalized expression data, requiring a different differential-expression framework from the GTEx analysis. Fourth, standard GTEx eQTL results are not sex-stratified, so sex-specific regulatory effects could not be directly assessed. Fifth, the hyperbolic embedding approach is heuristic rather than optimization-based, and alternative embedding methods may improve fit. Sixth, network geometry was assessed on largest connected components, which may not represent smaller components. Seventh, female-specific subnetworks were too small for robust geometric inference. Finally, both biomarker ranking schemes are composite and therefore depend on weighting choices; prospective validation in independent and sex-balanced cohorts is essential before clinical translation.

## 5 Conclusion

This multi-layer analysis shows that sex-dimorphic aging is a pervasive feature of CVD gene expression and that integrating expression, regulation, network geometry, genetic support and druggability substantially improves biomarker prioritization. The study identifies 35 high-confidence sex-dimorphic genes, highlights distinct geometric organization in complementary interaction networks and shows that most CVD genes have tissue-dependent regulatory architecture. It also nominates 19 actionable biomarkers supported by multiple evidence layers, approved drugs and GWAS colocalisation. Among these, *GUCY1A2*, *CACNA1D*, *PGR*, *PDE5A* and *LEPR* emerge as the strongest candidates for sex-stratified validation, with *GUCY1A2* and *PDE5A* converging on a nitric oxide–cGMP pathway that may contribute to sex-dependent vascular aging. These findings provide both a set of prioritized biological hypotheses and a general computational framework for developing sex-aware approaches to cardiovascular precision medicine.

## 6 Acknoledgements

The analysis were carried on Biomni platform.

